# A biological growth model using continued fraction of straight lines

**DOI:** 10.1101/2025.01.07.631841

**Authors:** Shruti I S, Vijay Prakash S

**Affiliations:** Independent Researchers, Alappuzha, Kerala, India

## Abstract

S-shaped curves are ubiquitous in biology especially when it comes to growth of a population or even an individual. Growth models such as the classical Verhulst-Pearl logistic growth equation and its extensions effectively model such S-shaped growth curves. Most of these models are parametrised by three or more parameters. In this work, continued fraction of straight lines has been applied to model S-shaped curves of biological growth through the use of only two parameters *a* and *m*. Here, *m* is the maximum growth rate and *a* is the parameter restricting the growth rate. The parameters *a* and *m* help to better interpret the data when compared to the logistic growth model since *m* represents factors promoting growth while *a* represents the constraints on growth. This model is effective for modeling both population as well as individual growth, especially around the phase of rapid growth.

## 1 Introduction

Biological growth can be studied in mainly two ways. One is physical growth of an individual or a group of individuals i.e. mean individual growth. The second way is the growth of a population i.e. increase in the number of individuals. In both cases, the growth follows a similar pattern [1]. This pattern when represented on an *xy* − plane gives an S-shaped curve also called as a sigmoidal or a logistic curve [2, 3]. This S-shaped graphical representation of biological growth can be interpreted in the following way. The initial phase of growth is gradual followed by a rapid increase known as the logistic phase. This rapid increase in growth continues until it reaches a maximum after which it again slows down until there is no observable change in growth any more [1]. Several models have been given to represent this growth pattern [3, 4, 5, 6]. In general, S-curves are modeled with logistic sigmoid function [3, 4, 5, 6] and the Verhulst-Pearl logistic growth equation is known to be the most successful one in representing restricted population growth. Further models are mostly extensions of this classical equation. In all these models, the exponential term is ubiquitous and parametrization is achieved with three or more parameters [7].

In this paper, we have presented a growth model based on continued fraction of straight lines parametrized with two parameters *a* and *m*. This model has been used to represent firstly, the population growth of yeast cells and *Drosophila* individuals grown as laboratory cultures [1]. The second attempt involves modeling mean growth of individual organisms, specifically of the sunflower (*Helianthus*) plant [8] and the male white rat [9]. We also model individual growth of a human male up to 18 years of age [10, 11].

In the following section, we describe continued fraction of straight lines with two parameters *a* and *m*. In sections 3, 4 and 5, we model population growth, mean individual growth and individual growth, respectively. We discuss our findings in section 6 and conclude our work in section 7.

## 2 Continued fraction of straight lines

Consider the parametric straight line equation *y* = *mx* with the parameter *m* as the slope on *xy* − plane. This equation is modified as

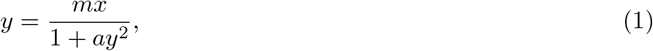

where *ay*^2^ bounds the values of *y* when *a >* 0. This modification with *a >* 0, behaves like a straight line as *y* → 0 but as *y* values increase with *x*, the line modifies in to a S-shaped curve. The real-valued solution of Eqn. (1) for *a >* 0 is given by

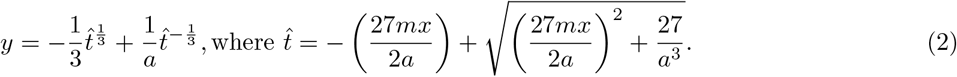

S-shaped curves of the above equation is provided in Fig (1). Eqn. (1) can be re-written as [12]

**Figure 1:**
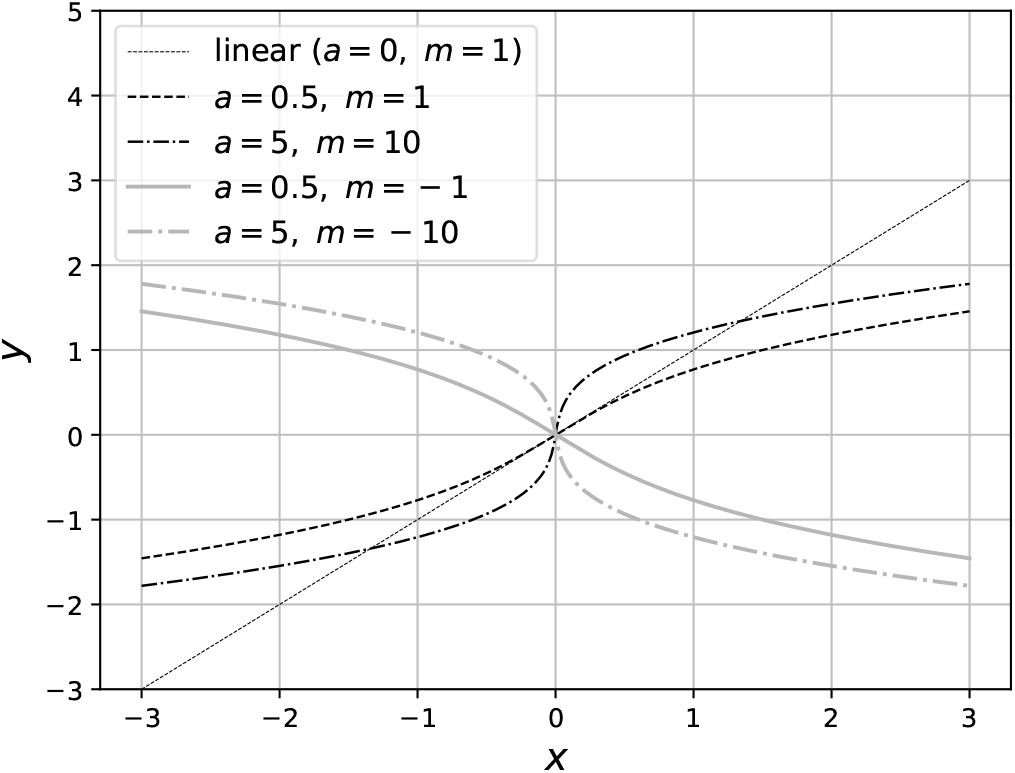
S-shaped curves for various *a* and *m*.

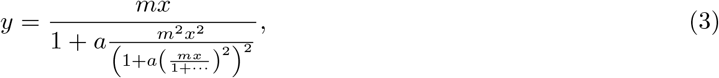

which is a continued fraction of *y* = *mx*. The above continued fraction converges to the solution of *y*, which is given by Eqn. (2). From Eqn. (3), it is known that *ay*^2^ in Eqn. (1) is a sum of series of bounding terms [12].

Thus, to model growth, we have presented a parametric equation which behaves like a straight line around the origin and takes an S-shape on *xy*− plane.

### 2.1 As a growth model

Differentiating Eqn. (1) we have

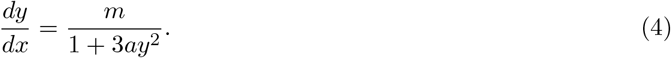

The maximum rate of change occurs at the origin i.e., *dy/dx* = *m* when *y* = 0 and it reduces as we deviate from the origin.

As a growth model, we consider the maximum rate of growth as the center or origin. Hence, Eqn. (1) can be written as

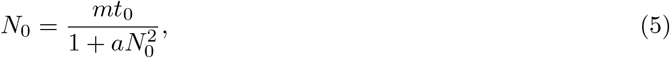

where *N*_0_ = *N* − *N*_*m*_ and *t*_0_ = *t* − *t*_*m*_. Here, *N*_*m*_ is the population size at time *t*_*m*_ of maximum growth rate *m*, with *N* as the actual population size and *t* as time. So the above equation can be re-written as

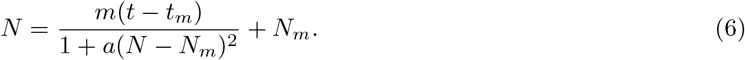

And for growth rate we have

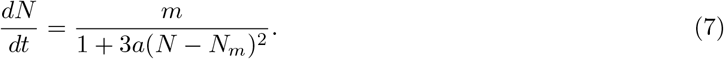

From the above, it can be seen that *m* represents the maximum growth rate at *N* = *N*_*m*_, while *a* acts as a restricting parameter that limits this linear growth. Thus, *m* is the growth promoting parameter because it represents the state when restrictions on growth become zero, i.e., *a*(*N* − *N*_*m*_)^2^ = 0 when *N* = *N*_*m*_. This leads to *a* being the restricting parameter of this maximum growth. Henceforth, we refer to Eqn .(6) as *a* − *m* model.

### 2.2 Comparison with the logistic model

According to the Verhulst-Pearl logistic growth model [2]

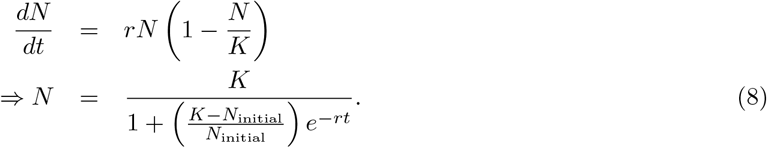

Here, *K* is the carrying capacity, *r* is the intrinsic growth rate and *N*_initial_ is the population size at initial time, *t* = 0. In this work, since we consider both the population and individual growth, we parameterize Eqn. (8) as

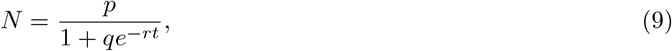

where *p, q* and *r* are the parameters [7, 13].

### 2.3 Fitting procedure

Both the models are fit using `lmfit` python package [14]. Initially, we define the models as separate functions

~~~
#Logistic model
def logistic(x,p,q,r):
  return p/(1+q*np.exp(-r*x))
# a-m model
  def am_model(x,a,m):
   t=(-27*x*m/2/a+np.sqrt((27*x*m/2/a)**2+27/a**3))
  return t**(-1./3)/a-t**(1./3)/3
~~~

We then use the `lmfit` to estimate the parameters *a, m, p, q* and *r* as follows.

~~~
# Create model from the above definitions
model_log = Model(logistic)
# Define positive parameters to be estimated with inital values
pars_log = model_log.make_params(p=dict(value=20,min=1e-9),
  q=dict(value=20,min=1e-9),
  r=dict(value=0.01,min=1e-9))
# Fit the model using least-squares minimization
out_log = model_log.fit(ydata,pars_log,x=xdata)
model_am = Model(am_model)
pars_am = model_am.make_params(a=dict(value=0.1, min=1e-9), m=dict(value=1))
#Here, xmod is xdata-mean(xdata) and ymod = ydata - mean(ydata)
out_am = model_am.fit(ymod,pars_am,x=xmod)
~~~

The goodness of fitting can be verified both graphically and by observing the `out.bic`, which is the Bayesian Information Criterion, since it is the most conservative measure for the goodness of fitting [14].

#### 2.3.1 Finding *N*_*m*_ and *t*_*m*_

In the following sections, both *a* − *m* model and the logistic model with parameters *p, q* and *r* are compared while fitting them for various growth data. We neglect some of the data points to get a good fit. Considering the selected data points, *N*_*m*_ and *t*_*m*_ will be the mean value of *N* and *t*, respectively.

## 3 Population growth

In this section, we fit the *a* − *m* model on the population growth data of *Drosophila* (Table 11 of [1]) and yeast (Table 9 of [1]). The three-parameter logistic model is compared with the two-parameter *a* − *m* model.

### 3.1 Drosophila

The growth model for *Drosophila* is shown in Fig. (2). In Fig. (2a), both the logistic model and the *a* − *m* model have been fitted for the entire data. In Fig. (2b), the first two data points have been removed. From Fig. (2a), it is observed that the logistic model fits better for the entire data compared to the *a* − *m* model. However, the *a* − *m* model fits well after simply omitting the initial two data points as shown in Fig. (2b). This fit made with two parameters is very close to the three-parameter logistic model. Thus, *a* − *m* model fits best for the phase that shows the maximum manifestation of population growth.

**Figure 2:**
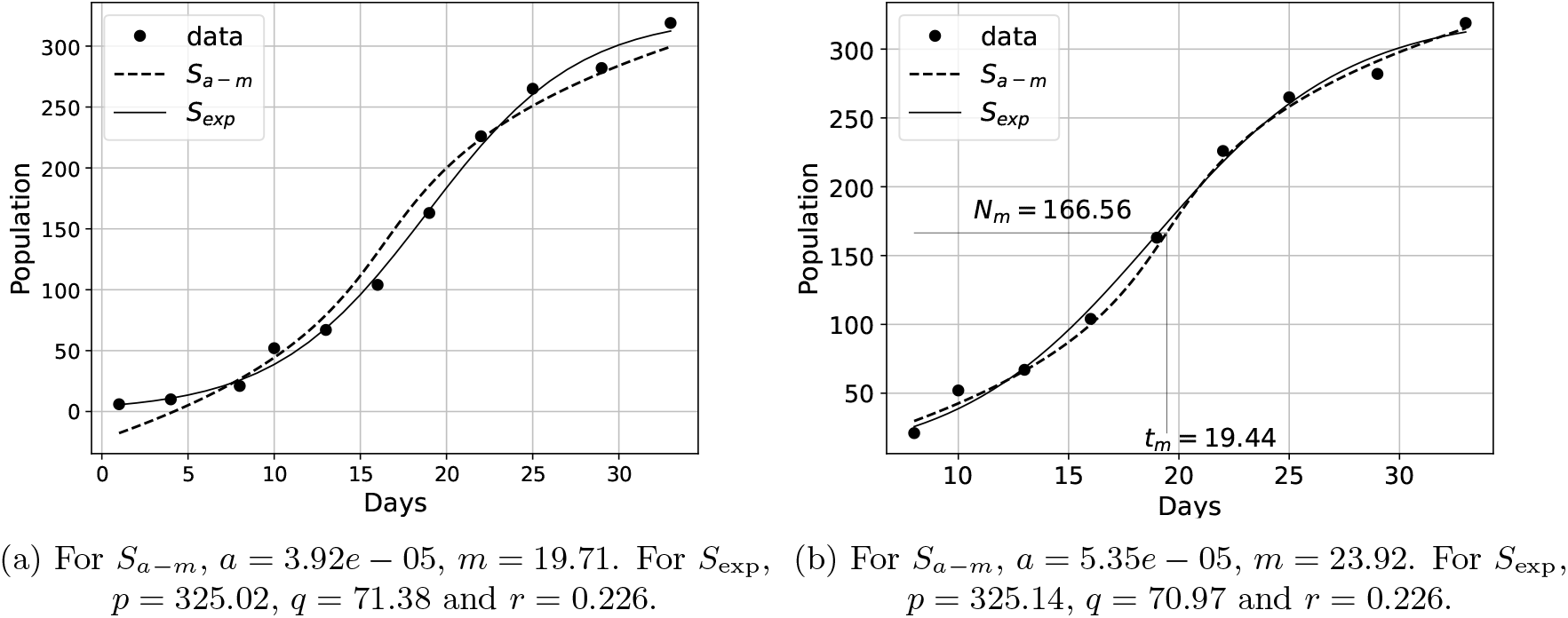
Comparison of the *a* − *m* model shown as *S*_*a*−*m*_ with the logistic model shown as *S*_exp_ for the *Drosophila* population data (Table 11 of [1]).

Based on Eqn.(6), the maximum growth rate is found to be *m* = 23.92. This means around 24 *Drosophila* individuals are born per day when the population size is 200 individuals i.e., when *N* = *N*_*m*_. As the population size deviates from *N*_*m*_, the growth rate decreases with the square of deviation which is parameterized with *a*.

### 3.2 Yeast cells

Here, we fit the *a* − *m* and logistic growth models on the yeast cell division data provided by Pearl [1] as shown in Fig(3). Similar to the Drosophila population model, the logistic model fits well for the entire data as compared to the *a* − *m* model as shown in Fig.(3a). However, around the rapid growth phase *a* − *m* model fits as well as the logistic one as shown in Fig. (3b). This is achieved by neglecting the first two and the last five data points. Also, a numerical value for the maximum growth rate is obtained directly from the parameter *m* which is around 103 yeast cells per day.

**Figure 3:**
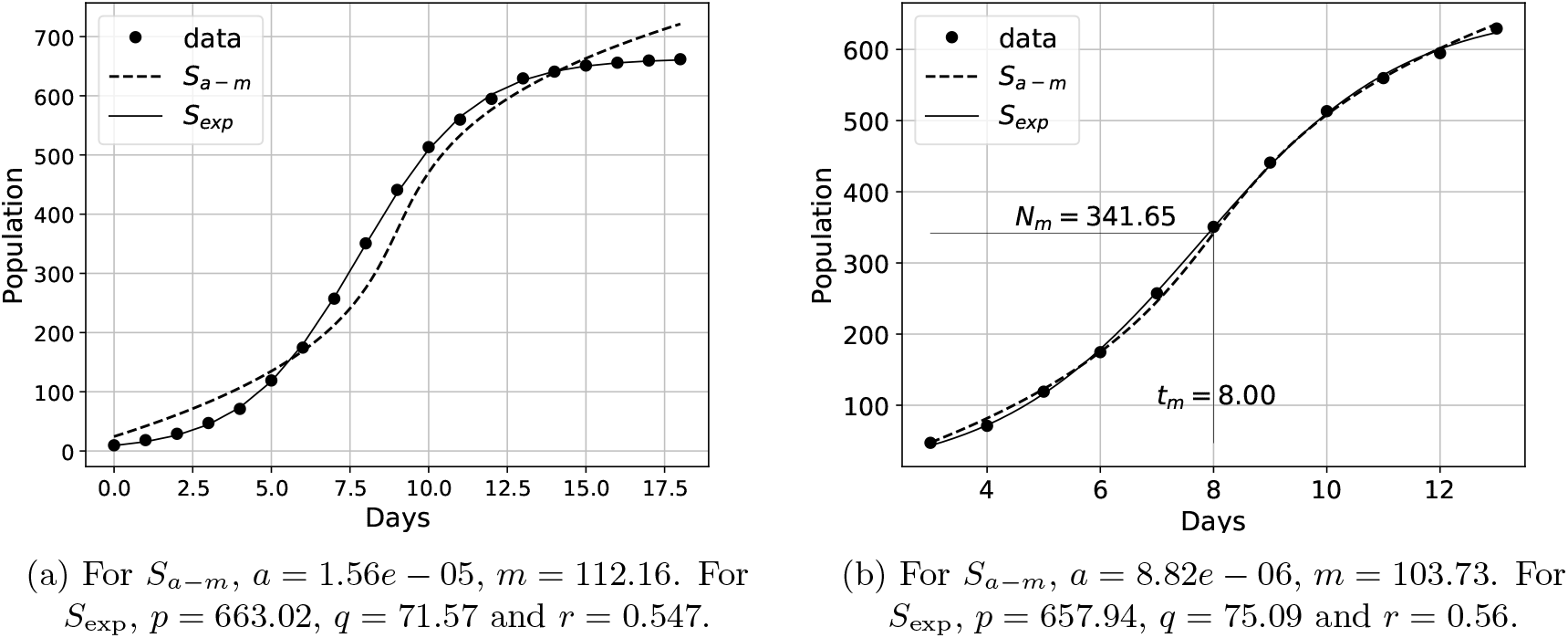
Comparison of the *a* − *m* model shown as *S*_*a*−*m*_ with the logistic model shown as *S*_exp_ for the yeast population data (Table 9 of [1]).

## 4 Mean individual growth

In this section and the subsequent section, we consider growth of individuals. This is done by simply replacing the population size *N* by a quantity *Q* under study. Here, we consider *Q* to be the height or weight of individuals. So, Eqn. (6) and Eqn. (9) are modified as

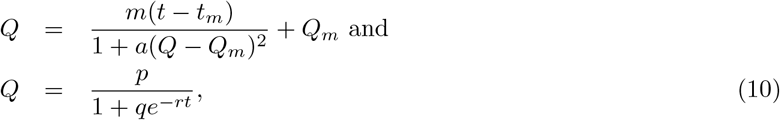

respectively. Here, *Q*_*m*_ is the value of the quantity under study when the rate of growth in *Q* is maximum, which occurs at time *t* = *t*_*m*_.

### 4.1 Mean height of sun flower plants

We now consider the mean height growth data of 58 sunflower plants (*Helianthus*) provided by Reed et al. [8]. Taking *Q* to be the mean height, we fit *a* − *m* model and the logistic model (Eqn. (10)). Similar to previous sections, the logistic model fits well for the entire data of mean height as compared to the *a* − *m* model as shown in Fig. (4a). However, after removing the final three points, *a* − *m* model fits well with the data as shown in Fig. (4b). Estimating *m* = 5.44 cm per day reveals that maximum growth rate of mean height is 5.44 cm in a day, when the mean height is around 133 cm.

**Figure 4:**
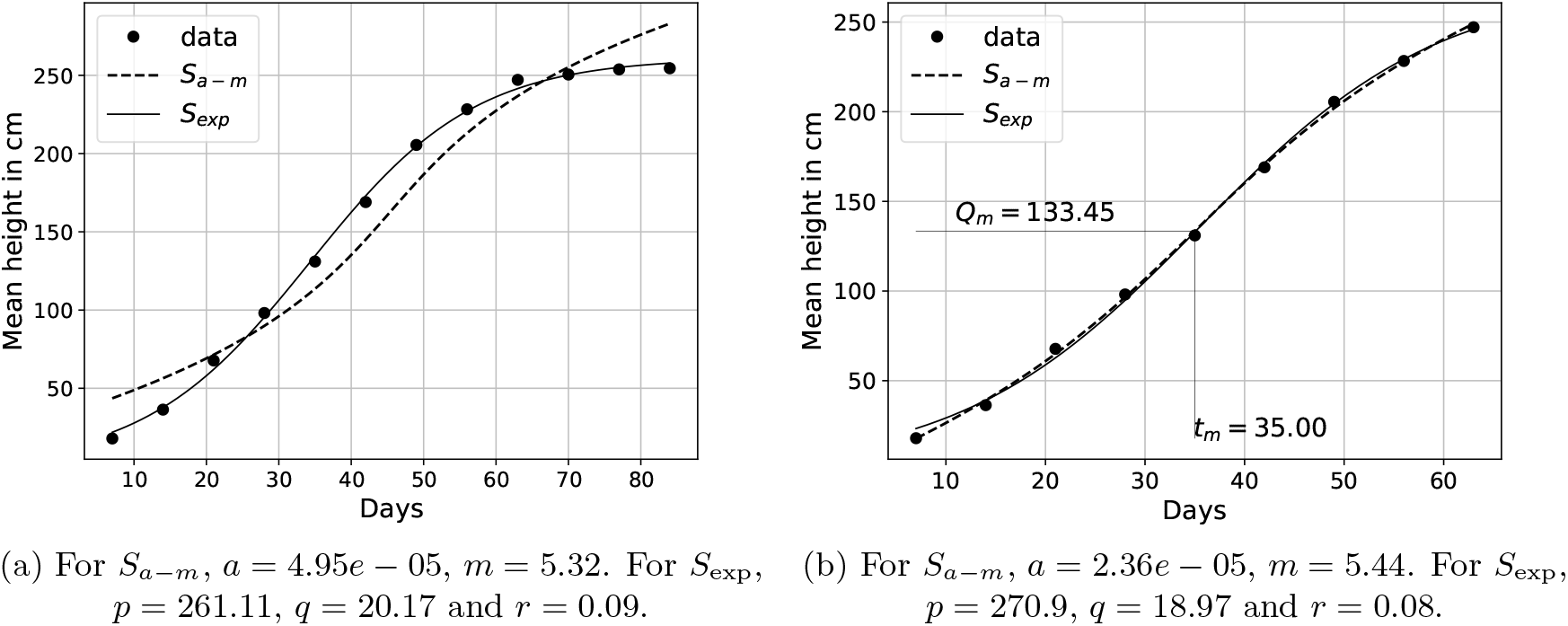
Comparison of the *a* − *m* model shown as *S*_*a*−*m*_ with the logistic model shown as *S*_exp_ for the growth in mean height of *Helianthus* plants [8].

### 4.2 Mean weight of male white rats

In this section we take *Q* to be mean weight in Eqn. (10) and consider the data of mean weight of male white rats provided by Donaldson [9] for fitting the growth models. Both the *a* − *m* model and the logistic model of Eqn. (10) do not fit well for the entire data as shown in Fig. (5a). However, after neglecting the final five points both the models fit well, especially around the phase of rapid growth as shown in Fig. (5b). *a* − *m* model fits even better when the initial ten points and the final eleven points are neglected as shown in Fig. (5c). In Fig. (5b) and (5c), we find that *m* = 2.36 − 2.63 grams per day, which is the maximum growth in weight of male white rats. Since Fig (5c) shows the best fit of the a-m model, it can be concluded that the estimation of mean values is more precise in this case.

**Figure 5:**
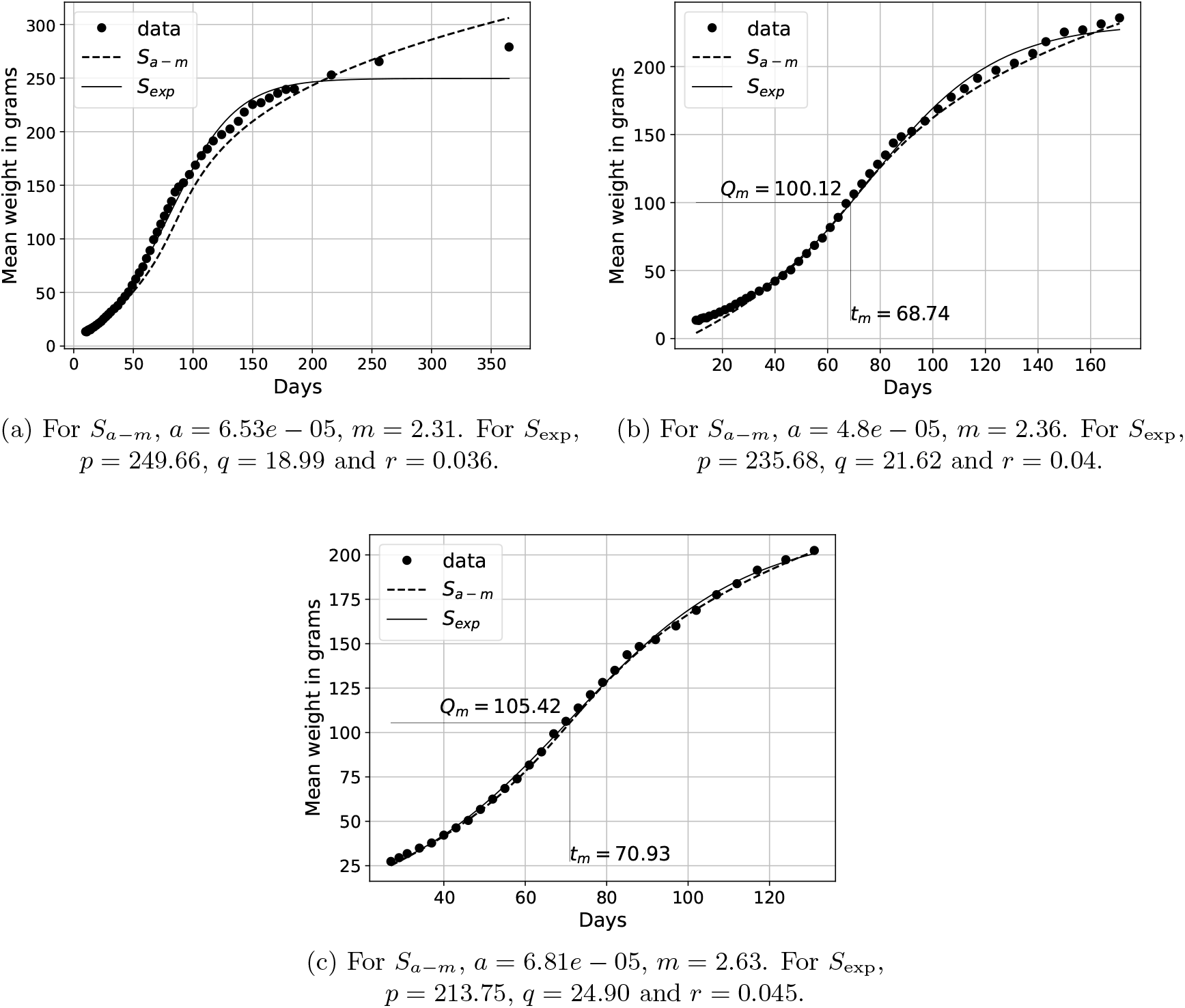
Comparison of the *a* − *m* model shown as *S*_*a*−*m*_ with the logistic model shown as *S*_exp_ for the growth in mean weight of male white-rats [1, 9].

## 5 Individual growth

So far, we have considered population growth data and the data of mean individual growth in height and weight of organisms. In this section, we consider the growth of an individual instead of a set of individuals. We consider the growth data provided as the first seriatum study of individual human growth [10, 11] over a period of 18 years. We initially try to fit both the models over the entire height data. As shown in Fig.(6a), both the models do not fit well over the entire data. Since we mainly look for S-shaped growth curve, we split the 18-year data in to smaller portions and obtain model parameters after fitting in these portions of data. In all these portions except one, the *a* − *m* model fits better than the three-parameter logistic model. The portions are as follows:

**Figure 6:**
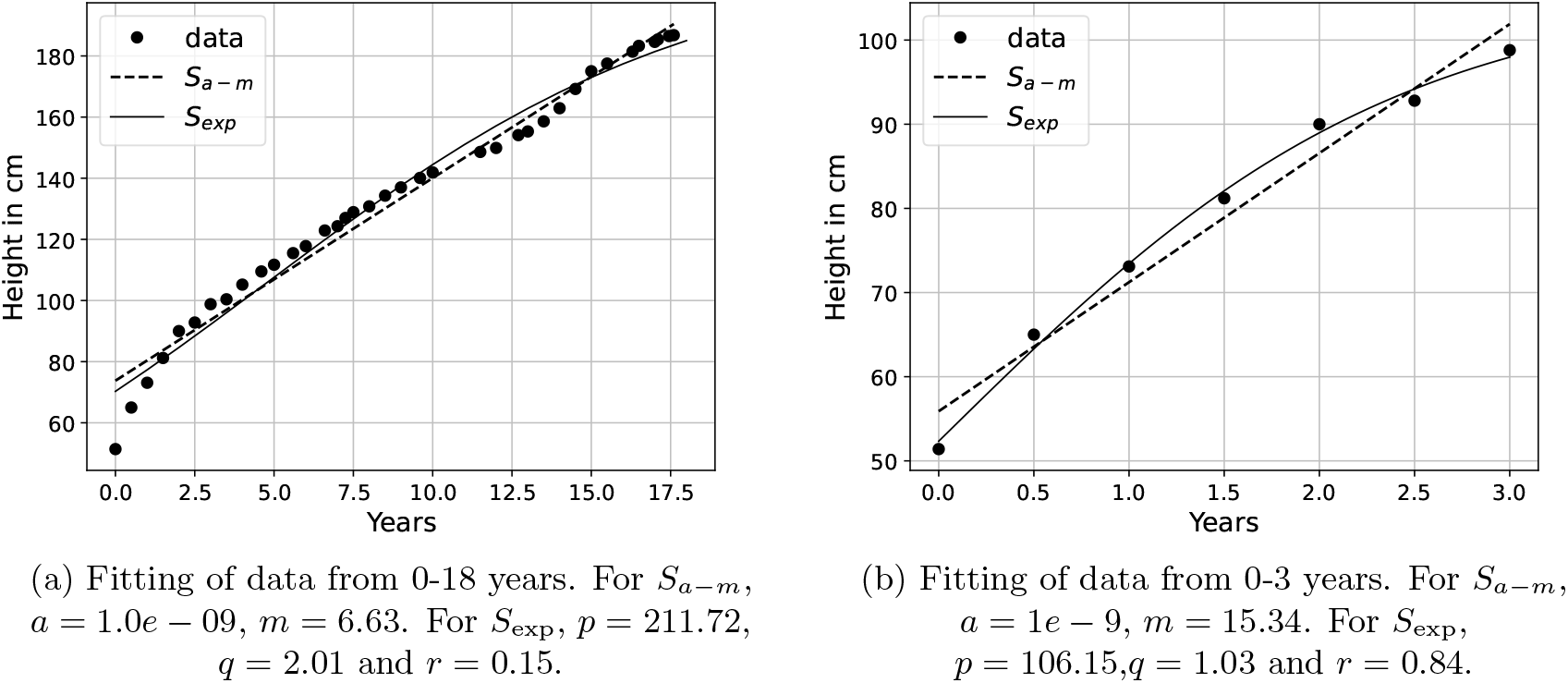
Comparison of the *a* − *m* model shown as *S*_*a*−*m*_ with the logistic model shown as *S*_exp_ for the growth in height of individual human male [10, 11].

- **0 to 3 years:** In this portion, the logistic model fits better than the *a* − *m* model as shown in Fig. (6b). This portion highlights the limitation of *a* − *m* model. We discuss the limitations in section 6.4.
- **3 to 5 years:** In this portion *a* − *m* model fits well as shown in Fig. (7a) and captures the nonlinearity better than the logistic model.
- **5.5 to 7 years:** The *a* − *m* model fits well for this portion of data in Fig. (7b) which is just four data points spread across two and a half years. Growth is more nonlinear, since this portion has the highest value of *a* compared to the rest of the fitted portions in the data.
- **7 to 10 years:** Both the models fit well for this portion of data as shown in Fig.(7c). Both the models show linear trend of growth. This portion has the lowest maximum growth rate which is *m* = 6cm per year.
- **11.5 to 17 years:** Perhaps this portion clearly shows that *a* − *m* model effectively captures the rapid phase of growth spanning many years of growth data as shown in Fig. (7d). This portion includes more number of years compared to the rest of the portions and the maximum growth rate is highest in this region, *m* = 11.89 cm per year.

**Figure 7:**
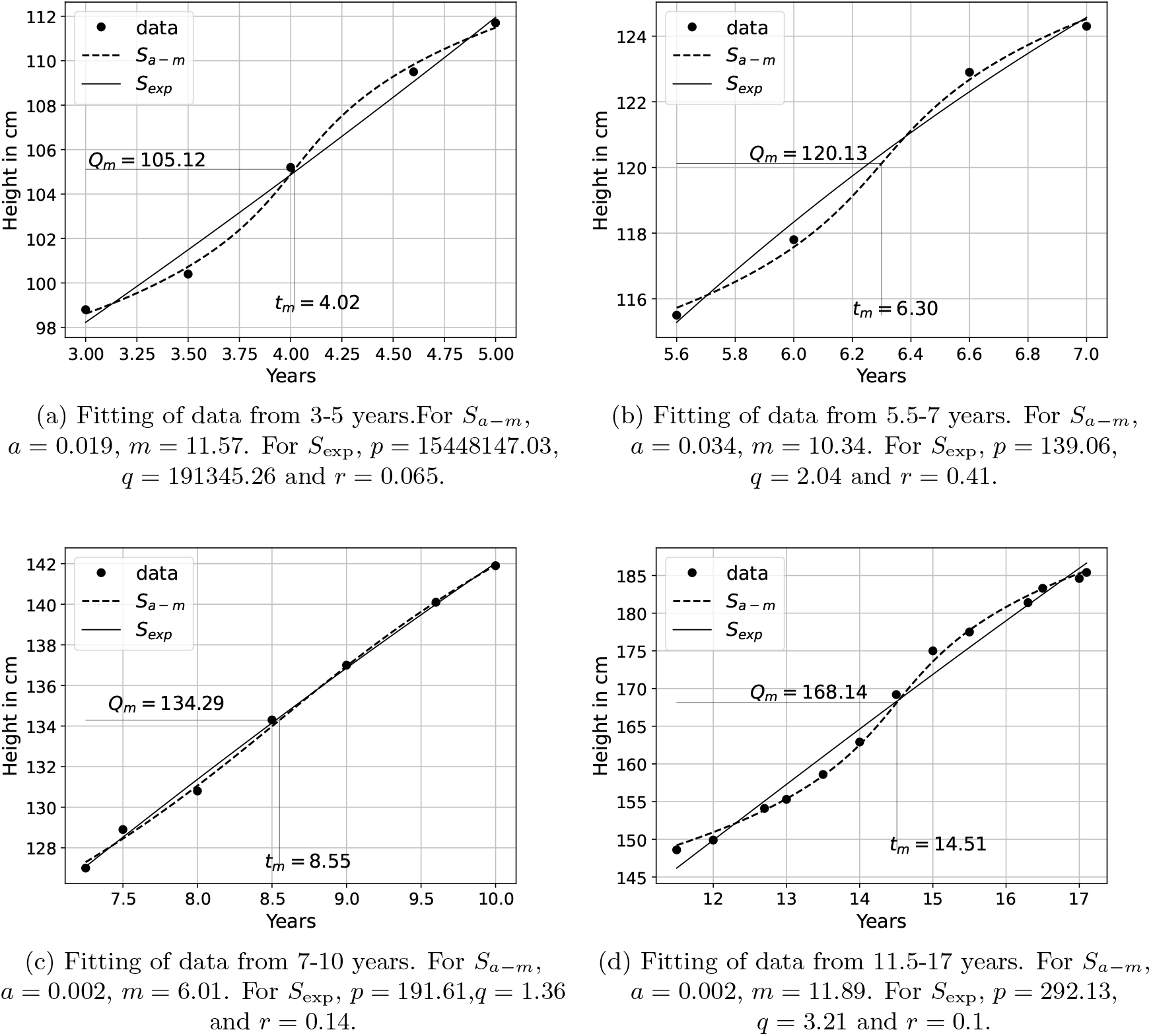
Comparison of the *a* − *m* model shown as *S*_*a*−*m*_ with the logistic model shown as *S*_exp_ for the growth in height of individual human male [10, 11].

## 6 Discussions

So far, in the previous sections we fitted the *a* − *m* model for various growth data and compared the fit with the logistic model. In this section, we present some of the possible discussions on biological growth based on the *a* − *m* model.

### 6.1 Growth is linear in time

Eqn. (1) can be re-written as

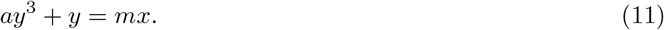

The term *ay*^3^ is a nonlinear parametric addition to the linear equation *y* = *mx*. When used as a growth model as seen in sections 3 and 4, it can be concluded that growth is fundamentally linear but restricted due to a nonlinear addition. This nonlinearity becomes more pronounced as we move away from the point of maximum growth rate.

### 6.2 Growth is symmetric around *t*_*m*_

From Fig. (1), it can be seen that the *a* − *m* curves vary symmetrically about the origin. Similarly, *a* − *m* growth model due to the term (*N* − *N*_*m*_)^2^ in Eqn. (6), varies symmetrically about *N*_*m*_ (or *Q*_*m*_). Since we have fitted the *a* − *m* model for population growth, mean individual and portions of individual growth, we can conclude that biological growth is symmetric about the point of maximum growth rate even if it is across many years as shown in Fig. (7d).

However, this symmetry is restricted to the parts closer to (*t*_*m*_, *N*_*m*_) or (*t*_*m*_, *Q*_*m*_). In other words, biological growth may not be completely symmetrical when all the data points are considered.

### 6.3 A growth coordinate system

Since we are using only two parameters to model growth, we can obtain planar plots with *a* and *m* as the axes. *m* − axis represents promoting factors for maximum growth and *a* − axis represents restricting factors on maximum growth. Thus, we can compare different growth data using this coordinate system as shown in Fig. (8). Since values of *a* vary over a wide range, it is convenient to use logarithmic scale representation for *a*− axis. From Fig. (8), it can be seen that growth of yeast (3) is more promoted and less restricted compared to the other population data. Similarly, human individual growth in height is more promoted and less restricted between 11.5 to 17 years compared to other portions of data in Sec. (5).

**Figure 8:**
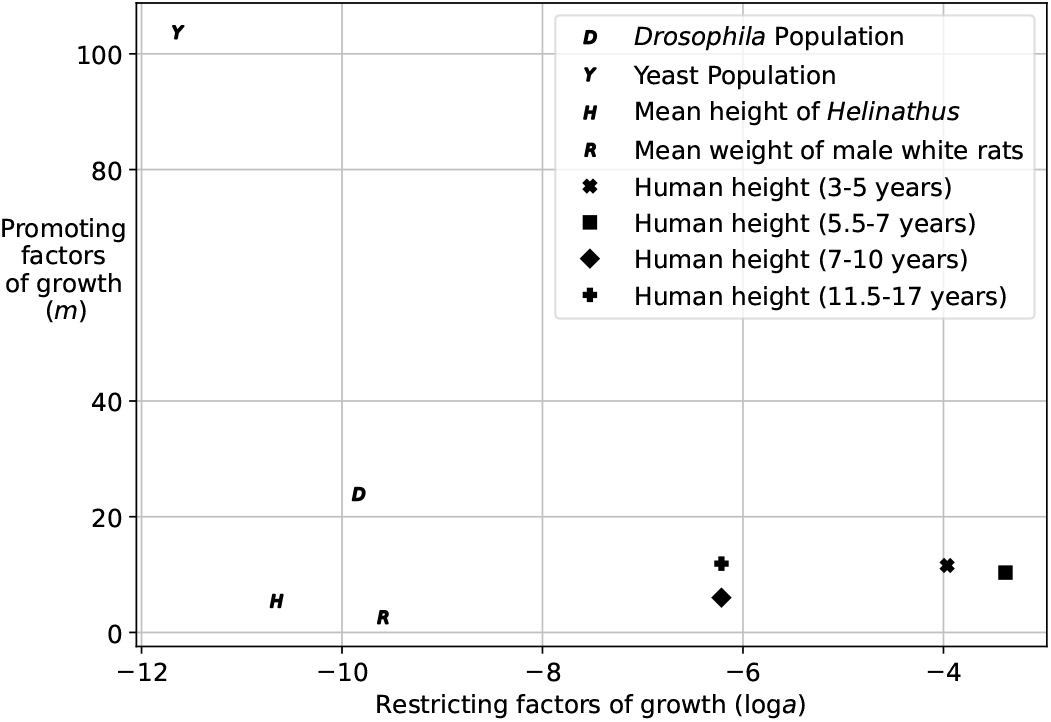
Growth coordinate system with log *a* as *x* − axis and *m* as *y* − axis. *a* and *m* values are from Figs. (2b),(3b),(4b),(5c),(7a),(7b),(7c) and (7d).

### 6.4 Limitations of *a* − *m* model

The *a* − *m* model fails to capture variations in the initial and final phases of population and mean individual growth. It also does not capture nonlinearities in Fig. (6b), which is the growth of an individual human male from 0-3 years. So, *a* − *m* model is mainly applicable to the parts of data around maximum growth rate. This also shows that the initial and final portions of population and mean individual growth data deviate from the (*N* − *N*_*m*_)^2^ nonlinearity, which in turn breaks the symmetry about (*t*_*m*_, *N*_*m*_) or (*t*_*m*_, *Q*_*m*_). This can be remedied in the future by adding more parametric nonlinear terms to Eqn. (1).

## 7 Conclusions

In this work, we have introduced a two-parametric growth model using continued fraction of straight lines. The two parameters are *a* and *m* which represent restricting and promoting factors of maximum growth, respectively. The *a* − *m* growth model is effective for modeling population growth, mean individual growth and to some extent, individual growth especially around the phase of rapid growth. This model suggests that biological growth is fundamentally linear in time but restricted by nonlinear factors. These nonlinear factors vary as the square of deviation from the point of maximum growth rate. Thus, the two parameters *a* and *m* can be considered as biological growth coordinates.

